# Mitigating activity mixing with personalized whole-brain modeling

**DOI:** 10.64898/2026.02.23.707368

**Authors:** Alina Suleimanova, Vladislav Myrov, Samanta Knapič, Wenya Liu, Paula Partanen, Maria Vesterinen, Satu Palva, J. Matias Palva

**Affiliations:** Department of Neuroscience and Biomedical Engineering, Aalto University, Espoo, Finland; Neuroscience Center, Helsinki Institute of Life Science, University of Helsinki, Helsinki, Finland; Unit of Psychology, Faculty of Education and Psychology, University of Oulu, Oulu, Finland; BioMag laboratory, Helsinki and Uusimaa Hospital Diagnostic Center, Helsinki, Finland; Centre for Cognitive Neuroimaging, School of Neuroscience and Psychology, University of Glasgow, Glasgow, UK

## Abstract

Mechanistic neuroimaging-based biomarkers based on localized brain activities or interactions are a central tenet in precision and personalized psychiatry. However, their accuracy may be limited by Activity Mixing that is a novel construct indicating the entanglement of any local neuronal activity with activities elsewhere in the network through long-range spatiotemporal correlations, which degrades the localization of brain–symptom associations. Here, we posit that the ill-posed inverse problem of activity mixing can be mitigated by fitting of generative whole-brain models. We developed a multi-objective fitting approach to estimate subject-specific local- and inter-areal brain-dynamics control parameters from neuroimaging observables. By integrating both synchronization and criticality metrics, this approach yields personalized parameters capture individual brain network dynamics more accurately than raw observables. *In silico* validation demonstrated that fitted model parameters improved brain–symptom correlation estimates by 30–85% and reduced false-negative rates by approximately ∼67% relative to conventional observables-based analyses. As *in vivo* proof-of-concept, resting-state magnetoencephalography (MEG) data from 230 patients with major depressive disorder (MDD) showed that aberrant brain criticality in the alpha-frequency band (11 Hz) was a significant predictor of disability with a correlation coefficient of 0.236 (95% confidence interval (CI) = [0.206, 0.266]) in 27 (CI = [23, 31]) significant cortical parcels. Model fitting both improved this correlation estimate by ∼56% up to 0.368 (CI = [0.341, 0.405]) and localized it ∼25% more narrowly to 20 (CI = [18, 22]) parcels. These findings suggest that model fitting can mitigate the effects of activity mixing and provide control-parameter estimates that delineate mechanistic biomarkers for brain disorders more accurately than the raw brain imaging observables.

## Introduction

Mechanistic biomarkers for mental disorders are a promising avenue to enable personalized or precision psychiatry solutions and constitute a pathway to refine the diagnostic processes for mental disorders^1^. Such biomarkers could comprise local brain areas or inter-areal interactions in well-delineated subnetworks exhibiting pathophysiological activity identifiable through correlations with clinical symptoms^2,3^. Functional connectivity (FC) derived from resting-state functional magnetic resonance imaging (fMRI) has been widely used to identify subnetwork alterations and symptom-specific network patterns in major depressive disorder (MDD)^4–7^, and mapping these brain–symptom associations can reveal clinically meaningful depression phenotypes^3,8^. Magnetoencephalography (MEG) and electroencephalography (EEG) extended this line of research by leveraging oscillation-based FC, which has enabled, *e.g*., identification of MDD subtypes^9,10^.

We posit here that “Activity Mixing” constitutes a central and currently poorly recognized limitation in the paradigm for biomarker identification. Activity mixing denotes the entanglement of any local neural activity with the activity dynamics of the network neighbors, leading to the mixing of activities in pathophysiological and putatively healthy brain areas and thus reducing their identifiability and separability. Notably, activity mixing refers to the mixing of the underlying neuronal activities per se, and is distinct from the mixing of the measurement signals, such as source leakage in electro- and magnetoencephalography^11,12^.

Several lines of evidence suggest that the brain operates near a critical transition between disordered (subcritical) and ordered (supercritical) phases of brain dynamics^13–15^, which exacerbates the activity mixing problem. The hallmarks of criticality include long-range temporal correlations (LRTCs)^13^, which predict behavioral variability^15^ and occur across a wide regime of critical-like dynamics extended by structural and functional heterogeneities^16–18^. Brain criticality is also evidenced by power-law scaled neuronal avalanches, observed across scales and species from rodents^19,20^ to humans^15,21,22^. Criticality has profound consequences for how local neural activities and interactions translate into population dynamics and observables.

In the subcritical phase, activity dynamics and inter-areal activity correlations in the form of phase-correlation-based functional connectivity, are largely determined by with the system’s underlying structure^23^, and hence activity mixing remains minimal. However, in systems operating at the critical phase transition, second-order interactions and spatio-temporal power-law correlations emerge, which drive a breakdown of structure–function coupling^23,24^ and lead to greatly increased activity mixing. Furthermore, there is large inter-individual variability in how close to peak criticality individual brains and brain systems operate in both healthy and brain disorder cohorts^14,16^, which may further drive inter-individual variability in the extent of activity mixing.

Despite the widespread use of MEG and EEG in clinical research, criticality-based metrics have remained rarely been used as biomarkers. Criticality provides a framework for a novel class of mechanistic biomarkers^25^ by linking cellular processes to emergent population-level dynamics. In particular, as the balance between excitation and inhibition (E/I) is likely the key control parameter for brain criticality^26^, metrics of criticality yield a window into E/I and its functional consequences, which are relevant across a wide range of brain disorders^25,27–29^. Altered LRTCs have been associated, *e.g*., with depression^30^, early-stage Alzheimer’s disease^27^, and epilepsy^29^. Thus, deviations in criticality observables across brain networks can act as disorder biomarkers reflecting how ‘hidden’ underlying synaptic- and cellular-level impairments manifest as observable abnormalities in system-level dynamics^13,25,30^.

We propose here a “reverse-engineering” approach to mitigate activity mixing by fitting the generative, whole-brain Hierarchical Kuramoto model with MRI and MEG data on structural and functional connectivity and brain criticality to infer subject-specific control parameters governing brain dynamics. Generative models based on the Kuramoto framework, including the canonical Kuramoto model^31^ and its extension into a multi-scale Hierarchical Kuramoto model^23^, simulate emergent collective synchronization dynamics, providing a mechanistic understanding of how small alterations in control parameters, such as coupling strength, can nonlinearly propagate through network interactions and reproduce large-scale brain dynamics observed in healthy brains and brain disorders^32,33^. In particular, the Hierarchical Kuramoto model enables modeling in-vivo-like criticality and connectivity concurrently as both facets of brain dynamics co-emerge in the model.ww

Parameters derived from personalized models, such as global coupling strength, have demonstrated clinical relevance in prior studies, for example in relation to depression severity^34^. However, previous studies in whole-brain model personalization have primarily focused on reproducing functional connectivity^35,36^ or metastable substate probabilities (PMS)^37,38^, rather than directly targeting the critical dynamics that shape large-scale neural organization^39^. Because critical brain dynamics may both be essential in underlying disorder mechanisms^27,28^ and contribute to activity mixing, we posit that incorporating criticality into model fitting is central for enabling the estimation of subject-specific control parameters.

Here we introduce and validate a model-fitting approach to mitigate activity mixing by integrating both criticality and synchronization metrics. We used the fitting of Hierarchical Kuramoto models to MEG-derived brain-dynamics observables constrained by subject-specific diffusion-weighted imaging (DWI)-derived structural connectomes to estimate personalized parameters governing local and inter-areal coupling and criticality. The fitted model parameters provide mechanistic insight into the drivers of system-level dynamics and deviations from criticality. We hypothesize model fitting can mitigate activity mixing with improved brain–symptom correlation estimates using subject-specific model parameters.

## Results

### Activity mixing confounds mechanistic brain-symptom relationships

In complex systems such as the brain under conditions of criticality, signal propagation leads to activity mixing among nodes, complicating the identification of the underlying mechanisms that govern brain functions. To illustrate this concept, we present a hypothetical system (Figure 1a), where the mechanism generating the activity exhibits a perfect correlation with an external feature, characterized by a correlation coefficient of 1 (Figure 1b). However, at the observational level, whether the data can be empirical or a result of generative forward modeling, activity mixing blends signals from different nodes (Figure 1c). This mixing leads to a reduced correlation at the nodes within ground-truth (GT) mechanism subsystem that are directly involved in processing the external feature (Figure 1d). Simultaneously, this mixing can result in an increase in correlations at neighboring nodes (see Figure 1d), which can complicate interpretations of the underlying mechanism.

**Figure 1.**
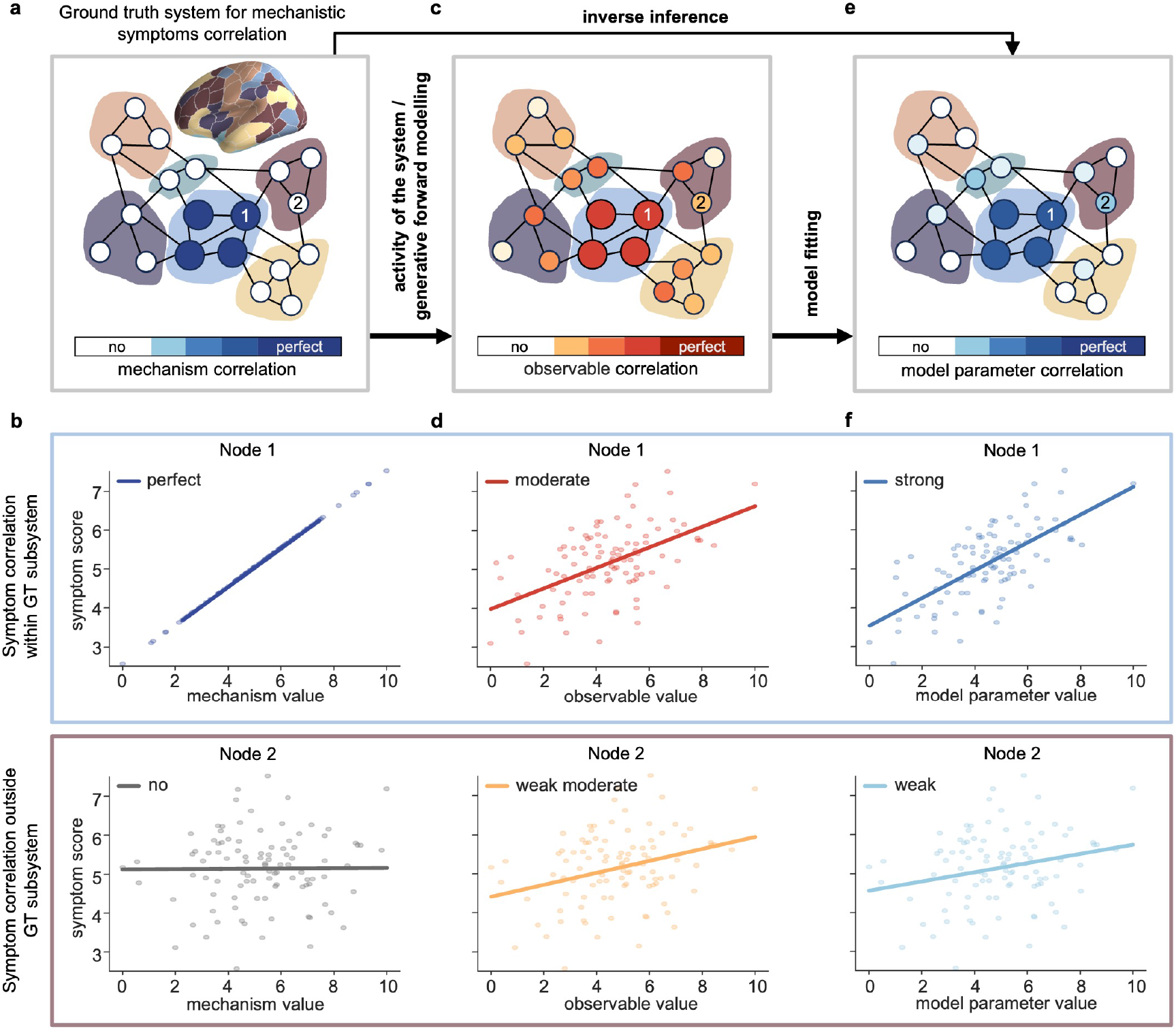
Schematic of activity mixing and its mitigation with model fitting. **a** Hypothetical, universal ground-truth (GT) system with a “mechanism” that is correlated perfectly with an external feature. **b** Correlation coefficients of GT mechanism with an external feature at target (blue) and non target (gray) nodes. **c** At the observational level, the system’s activity correlates spread to neighboring nodes. **d** Correlation coefficients of observables of a system with an external feature at target (red) and neighboring (yellow) nodes. **e** Reverse engineering with model fitting would reduce mixing and yield a mechanistic network estimate that is closer to GT. **f** Correlation coefficients of restored parameter of a system using fitting with an external feature at target (blue) and neighboring (light blue) nodes.

To mitigate the challenges posed by activity mixing, we propose employing reverse engineering techniques using model fitting. These approaches aim to reduce mixing, providing a network estimate that is closer to the actual system (Figure 1e). After fitting the model, the correlation coefficients at the target nodes are supposed to increase, indicating a clearer relationship with the external feature. In contrast, the correlations at the neighboring nodes decrease, helping to clarify the specific contributions of the target nodes to the overall dynamics (Figure 1f).

### Parameter estimation using personalized model fitting

We developed a novel algorithm that implements our approach to mitigate activity mixing by incorporating both functional connectivity and criticality properties in the estimation of model parameters (Figure 2a–c). To achieve this, we utilized the gradient direction as a function of a control parameter, approximating the synchrony metric, Phase-Locking Value (PLV), as a sigmoid function, and the criticality metric, Detrended Fluctuation Analysis (DFA), as a Lorentzian function due to its prominent peak and long tails.(see Figure 2c).

**Figure 2.**
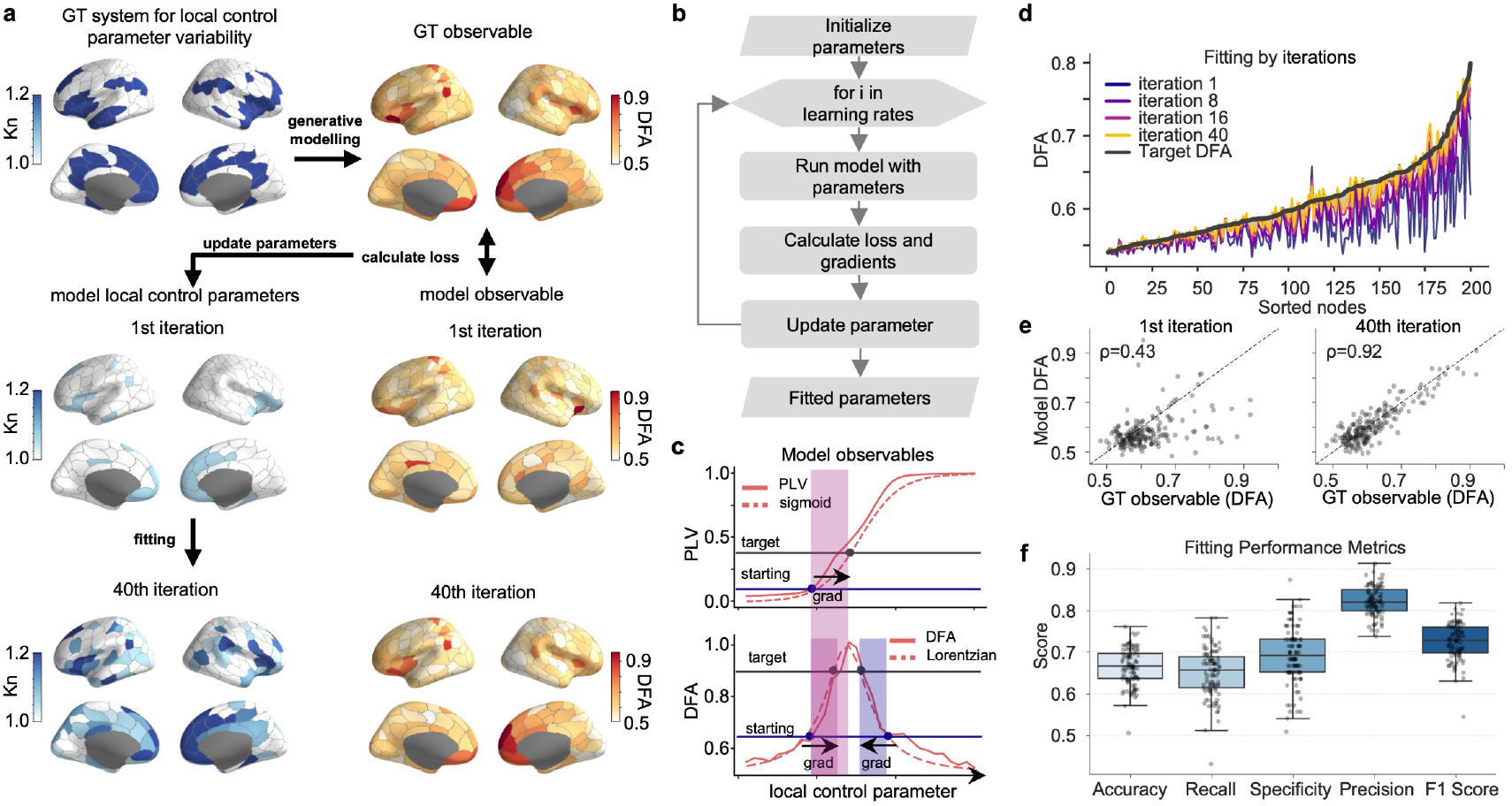
Model fitting approach and its evaluation. **a** An example of model fitting. An artificial phenotype created by multiplying the local control parameters Kn at selected nodes by 1.2. Learned parameters of a model fitted to a predefined phenotype. **b** A flowchart of the fitting algorithm. **c** The gradient direction as a function of a control parameter. **d** Changes in loss between target and fitted observables over fitting steps. **e** Scatterplots of a phenotype and a fitted model observables for the first and for the last step. **f** Performance metrics of the fitting method.

To validate the algorithm, we first generated artificial phenotypes with known changes in the control parameters and fitted a model to them. We created a vector of phenotype coefficients by increasing local control parameters in selected functional zones (Figure 2a). Then, we ran a phenotype-based simulation, calculated the PLV and DFA observables, and used them as target metrics in fitting.

We found that after fitting, the model was able to reproduce the altered dynamics with high similarity, with the average Pearson correlation coefficient across 100 subjects being 0.83 for DFA and 0.9 for PLV (Supplementary Figure 1). The scatters illustrate the similarities between the observables of the target and the fitted model, demonstrating the effect of the fitting by comparing the first and last steps (Figure 2e), while Figure 2d further shows that the loss between the target and the fitted DFA decreased progressively over these steps reflecting the fitting progress.

To validate the effectiveness of the developed fitting method, we calculated the performance metrics (Figure 2f) by converting the phenotype and the fitted parameters into a binary profile, where 0 represents no change and 1 indicates a change in the parameters. For *N* = 100, the fitting results achieved an accuracy of 0.66, indicating that it correctly classified 66% of the instances. It showed a recall of 0.65, capturing 65% of true changes, and a specificity of 0.82, correctly identifying 82% of cases without change, while the F1 score of 0.72 reflects a good balance between precision and recall. Moreover, the fitted parameters also followed an anatomical pattern similar to the GT values (see Figure 2a).

### *In silico* validation of model fitting approach

Brain disorders such as MDD are often heterogeneous and can affect different neural subsystems across individuals. To reflect this variability, we considered a scenario where local coupling in a cortical system constituted a mechanism for a hypothetical brain disorder. We employed five distinct mechanisms in our model of diseased activity, each corresponding to a functional subsystem (Figure 3a).

**Figure 3.**
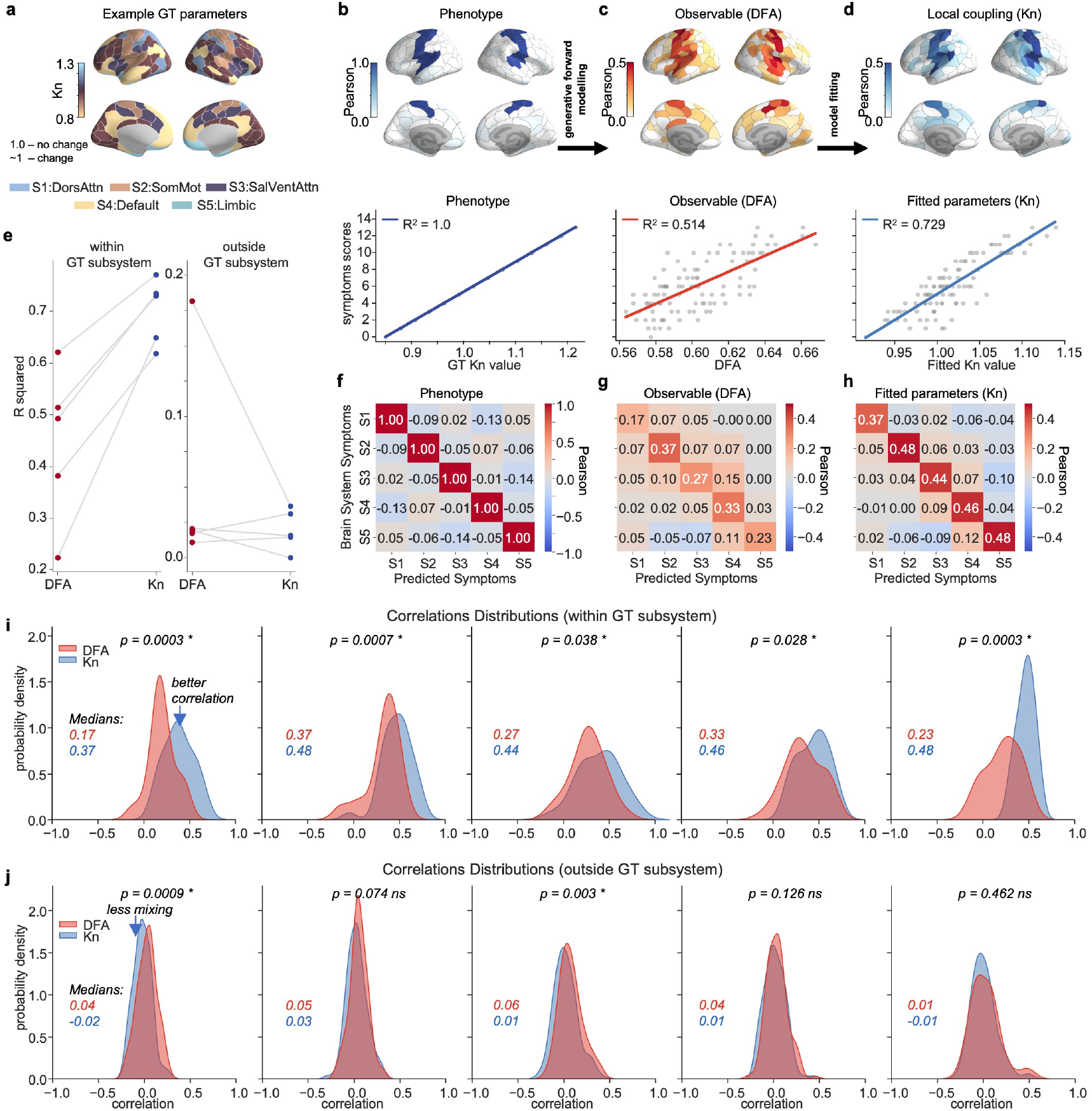
*In silico* proof-of-concept for mitigating activity mixing using model fitting. **a** A phenotype where parameters of five different functional networks (Dorsal Attention, Somatomotor, Salience and Ventral Attention, Default Mode, and Limbic) altered from default parameter 1. **b** Local coupling in the sensorimotor cortex perfectly correlates with a hypothetical clinical or behavioral feature (e.g., symptom severity) with Pearson’s correlation coefficient = 1 in *N* = 100 virtual patients. **c** Activity mixing reduces nodal criticality correlations, even with perfect observation. **d** Model fitting mitigates activity mixing, restores correlations and, reduces anatomical spread. **e** Correlation between the average values of observables and parameters with symptom severity within target and non-target subsystems. **f-h** Confusion matrices showing median correlations of subsystem parameters, observables, and fitted parameters with symptoms in *N* = 100 virtual patients. Distributions of diagonal elements are shown in panel i, and distributions of all off-diagonal elements for one subsystem are shown in panel j. **i** Distributions of parcel-level Pearson correlations within GT subsystems (diagonal values in confusion matrices in f-h). Model parameters correlated more strongly with symptoms than observables in target areas (two-tailed t-test, *p <* 0.05). **j** Distributions of parcel-level Pearson correlations outside GT subsystems (non-diagonal values in f-h) of observables and parameters.

We set parameters such that the GT local coupling parameter exhibited a perfect correlation (correlation coefficient = 1) with a hypothetical clinical or behavioral feature (e.g., symptom severity) in *N* = 100 virtual patients, as shown for the sensorimotor cortex subsystem in Figure 3b. However, despite this perfect correlation, the correlations of nodal criticality (DFA) observable with symptom severity were diminished due to activity mixing within the model (Figure 3c). To address the activity mixing issue, we applied the model fitting approach. Using linear regression, we found that compared to the correlations obtained from observables (*R*^2^ = 0.514) the restored local coupling parameters achieved 42% higher correlations (*R*^2^ = 0.729) within the sensorimotor cortex subsystem, while reducing anatomical spread effectively mitigating the effects of activity mixing (Figure 3d).

We further analyzed the relationship between the average values of the observables and the parameters relative to the symptom severity, both within GT subsystems and outside of them (Figure 3e). We found a 30-85% increase in the correlation coefficient with the fitted parameters compared to observables within the GT subsystem.

Then, to illustrate the median correlation values between symptom severity and subsystem-level parameters or observables, we created confusion matrices for the pairwise correlation coefficient with symptom severity. The GT local coupling parameter indicated a perfect correlation with the severity of symptoms (Figure 3f). The confusion matrix for the observables (Figure 3g) revealed that activity mixing reduced correlations within the GT subsystems, while the fitted local coupling parameters demonstrated stronger correlations within these subsystems (Figure 3h).

We also performed an analysis of parcel-level correlation distributions, which highlighted differences in correlation distributions between observables and fitted model parameters. Within the GT subsystems, the model parameters showed a significantly stronger correlation with the severity of symptoms compared to the observables demonstrating the effectiveness of the fitting approach in mechanisms discovery (Figure 3i). Outside of the GT subsystems, two of the five subsystems exhibited a significant difference (two-tailed t-test, *p <* 0.05) in the correlation distribution between the parameters and the observables (Figure 3j). Moreover, the model parameters exhibited a ∼67% lower false negative rate, further underscoring its sensitivity to underlying symptom-related mechanisms (Supplementary Figure 2).

### *In vivo* proof-of-concept biomarker identification

Depression symptoms exhibit broad variability across individuals, suggesting that symptom severity may relate to alterations in neural dynamics. To examine this, we focused on the relationship between criticality, quantified using DFA, and symptom severity measured by the Sheehan Disability Scale (SDS)^40^ in a resting-state MEG dataset of 230 patients with MDD. We targeted the alpha band at 11 Hz, where DFA values demonstrated the strongest association with the SDS symptom severity (Figure 4a).

**Figure 4.**
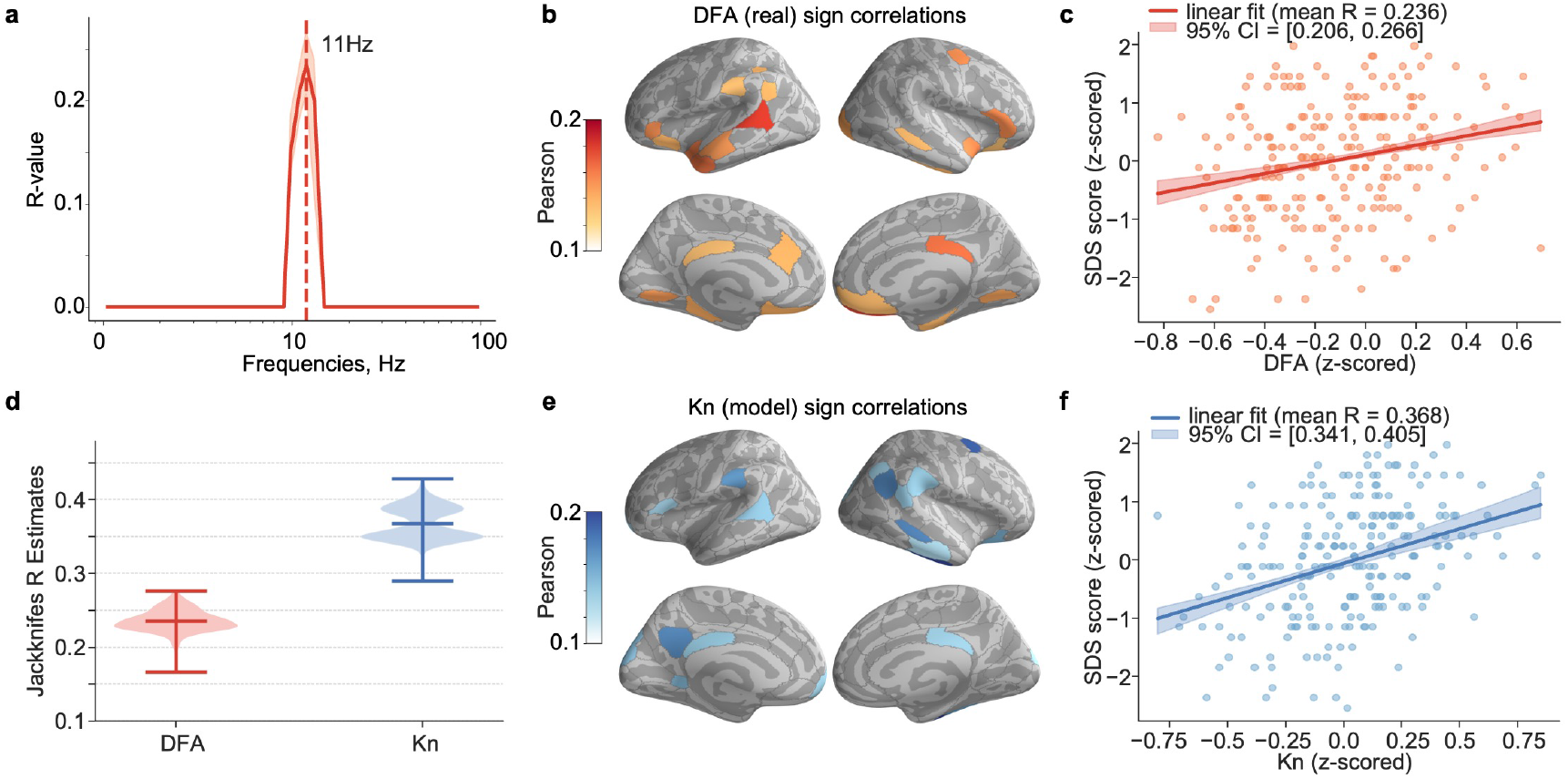
*In vivo* proof-of-concept for phenotyping using model fitting. **a** Spectral distribution of R-values of linear regression fit (confidence intervals (CIs) were calculated using Jackknifes estimates) for patient criticality observables (estimated using DFA) significantly correlated with Sheehan Disability Scale (SDS) symptom score. **b** Anatomical map of significant Pearson correlation coefficients between DFA and SDS symptom across brain parcels at 11Hz. **c** Scatter plot showing the relationship between z-scored SDS symptom values and average of z-scored DFA values from significantly correlated parcels, with linear regression fit of these values. **d** Jackknife estimates of R-value for nodes and parcels significantly correlated with SDS symptoms, for both local coupling and DFA. **e** Anatomical map of significant Pearson correlation coefficients between fitted local coupling parameters and SDS symptom across model nodes at 11Hz. **f** Scatter plot of z-scored SDS symptom values versus average of z-scored values of significantly correlated nodes of local coupling parameters, with linear regression of these values.

Then we analyzed Pearson’s correlation values between DFA and SDS symptom severity across brain parcels at 11 Hz, finding that the correlation spread across a wide range of brain regions (Figure 4b) including Default Mode, Control, Limbic and Visual Networks. However, the mean R-value of the average of the z-scored values of significant parcels was modest and equal to 0.236 (Figure 4c).

Jackknife resampling revealed that the distributions of the R-values for the fitted parameters and the MEG observables were essentially non-overlapping (Figure 4d). The jackknife R-values for the correlations between the fitted parameters and SDS symptoms (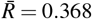, 95% confidence interval (CI) = [0.341, 0.405]; Figure 4f) were ∼56% stronger than the correlations between the MEG observables and SDS symptoms (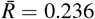, 95% CI = [0.206, 0.266]; Figure 4c; see Figure 4d).The difference was significant (two-tailed t-test, *p* = 1.32 × 10^−258^), effectively approaching zero. A bootstrap analysis yielded consistent results, again showing that the fitted-parameter estimates were significantly higher than the MEG observables (Supplementary Figure 3).

When exploring the relationship between SDS symptoms and the fitted local coupling parameters, we found that the same functional subsystems were involved as those identified by DFA of MEG data (Figure 4e). Furthermore, we evaluated the correlations of the DFA of the fitted model with the symptoms of SDS against the correlations with MEG data, with a Jaccard index of 0.52 (Supplementary Figure 4a) indicating a significant similarity compared to surrogates in the profiles between the model and MEG observables (see Figure 4b, Supplementary Figure 4b).

## Discussion

We posit here that empirical correlations between brain activity and clinical or behavioral observations may be confounded by a novel concept of “activity mixing”. Whereas the widely acknowledged “signal mixing” or “source leakage” result from the mixing of signals multiple underlying sources, especially in electrophysiological measurements, activity mixing arises when local neural dynamics *per se* become entangled with global network fluctuations. We show here that activity mixing even under conditions of zero signal mixing leads to an underestimation of true brain-symptom correlations and an overestimation of spurious ones. We hypothesized that fitting of observational data with a generative whole-brain model would mitigate activity mixing and allow brain-symptom relationships to be estimated through optimized model parameters. *In silico* validation confirmed that model fitting restored stronger and anatomically more specific associations between model parameters and symptoms. Because criticality amplifies the effects of activity mixing, we incorporated both synchronization and criticality metrics into a multi-objective gradient fitting approach. As an *in vivo* proof-of-concept, we applied our approach to MEG data from *N* = 230 patients with MDD, where model parameters showed significantly stronger and more localized correlations with SDS symptom scores than the conventional DFA observables. These results demonstrate that parameters derived from personalized brain models yield more accurate brain-symptom correlations than conventional observational approaches.

Activity mixing, where observed activity at a node reflects both local dynamics and propagated influences from connected nodes, has been documented for diffusion^41^ and contagion^42^ processes, as well as for information flows^43^ where activity mixing blurs the origin of activity. Brain networks are predisposed to exhibit activity mixing because their hierarchically modular architecture linked by hubs and intermodular pathways^44,45^ enables any local activity and inter-areal functional connectivity to be influenced by indirect or higher-order pathways rather than direct local interactions^46,47^. Moreover, because brains *in vivo* are thought to operate within the critical regime^19,48–50^, even small perturbations can spread widely through hub connections, blurring the distinction between local and distributed dynamics. Crucially, proximity to criticality^16,51^ modulates the degree of activity mixing and determines the extent to which local measurements are shaped by propagated signals. To address these challenges, we implemented the criticality-informed fitting approach. By incorporating critical dynamics, our approach effectively mitigates activity mixing, yielding stronger and more localized correlations between model parameters and symptom severity, as confirmed *in silico* validation (see Figure 3f).

Fitting methods are essential tools for solving the inverse problem in brain modeling, where the primary objective is to infer model parameters that can reproduce observed neural dynamics^52,53^. Fitting is commonly performed by optimizing models to reproduce neuroimaging data, such as FC, and often relies on large-scale simulations of Blood Oxygenation Level-Dependent (BOLD) signals^35^. In addition, recently, whole-brain models have also been used to estimate metastable substate probabilities (PMS)^54^, which are interpreted as indirect markers of critical brain dynamics^37,38^.

Despite recent advances, most existing fitting approaches rely on indirect markers of criticality. Gradient-based fitting, while effective in reproducing empirical covariance and functional connectivity patterns^52,54^, has been applied largely without explicitly targeting critical dynamics. However, criticality is essential for interpreting large-scale neural organization^39^. Our method introduces a novel fitting approach that integrates criticality metrics into the fitting process, allowing the model to capture both connectivity patterns and individualized signatures of critical brain dynamics. The method builds on the dependence of dynamics as function of the parameters of the Hierarchical Kuramoto model^23^ which enables efficient gradient-based fitting and faster convergence (see Figure 2d,e) than computationally intensive Bayesian methods^55,56^.

As an *in vivo* proof-of-concept, we applied our approach to resting-state MEG data from *N* = 230 patients with MDD. Discovering reliable neuroimaging biomarkers for mental disorders, such as depression, remains challenging due to the heterogeneity of these disorders^57,58^ and the confounding effects of activity mixing. Depression is associated with widespread structural and functional changes^59^, and functional MRI and MEG/EEG studies have reported alterations in FC, including default mode, control, and limbic networks^4,59,60^. Beyond FC, features such as LRTCs in the alpha and theta bands have been linked to the severity of depression^13,30^ and are thought to reflect shifts in the brain’s operating point away from criticality^16,51^.

Conventional correlation analysis between DFA values and SDS scores revealed spatially widespread associations spanning default mode (DMN), control, limbic, and visual networks (see Figure 4b). The involvement of DMN is consistent with prior studies of its altered activity and connectivity in depression^4,60^, but the modest correlation strengths suggested that these associations may be diluted by activity mixing^46^. Following model fitting, the resulting model coupling parameters showed stronger correlations with symptom severity, retaining prominent involvement of the DMN, control, and limbic networks. This improvement demonstrates that model-based metrics can enhance the identification of reliable biomarkers and is in line with previous evidence, where Kuramoto-derived coupling parameters were associated with depression severity^34^.

Recent studies have used whole-brain models to simulate neural phenomena, including cognitive state transitions^61,62^, responses to stimulation^37,38,63^, and the effects of structural lesions^36,64,65^. Extending this line of work, we applied the Hierarchical Kuramoto model^23^ to capture the synchronization patterns and criticality dynamics observed in MEG data from patients with depression. The Kuramoto model with incorporated EEG-derived connectivity has also been proposed to identify patient-specific brain stimulation targets that optimize functional network recovery^66^. By personalizing the model with individual diffusion-weighted imaging (DWI)-derived structural connectomes, we ensured that the model captured subject-specific brain organization. Similarly, personalized models constructed with diffusion tensor imaging (DTI) and MEG data have shown improved predictive power in clinical manifestations of conditions such as multiple sclerosis^67^. In epilepsy, personalized models constructed using DTI data have been used to simulate seizure propagation, improving the precision of surgical planning^68,69^. Comparable approaches have also been applied to stroke, where whole-brain models predict changes in functional connectivity^64^ and criticality^65^. These insights underscore the potential of integrating structural and functional data within generative models to advance the mechanistic understanding of brain disorders. Here, we introduce an approach for personalized generative modeling, in which models built with subject-specific structural connectomes are fitted to criticality and synchronization dynamics. The resulting personalized parameters can be used for biomarker discovery, with reduced activity mixing.

In conclusion, the present study establishes a proof-of-concept for criticality-informed personalized whole-brain modeling and highlights the potential of such model fitting in mitigating the activity-mixing problem.

## Methods and Materials

Hierarchical Kuramoto-based generative brain dynamics models were implemented to simulate critical dynamics using individual DWI-derived structural connectomes as weight matrices^16,23^. The model parameters were optimized with developed multi-objective gradient descent algorithm to reproduce individual-level MEG synchronization and criticality metrics.

LRTCs were quantified using DFA^70^, computed in the Fourier domain for computational efficiency^71^. DFA exponents were estimated by fitting fluctuation functions across window sizes with robust linear regression using a bisquare weight function^72^.

After fitting the model, the resulting fitted parameters were evaluated for their potential to capture individual diagnostic information beyond that provided by the MEG data alone. Pearson correlation analyzes were performed between clinical symptoms scores, model parameters, and MEG observables.

### Data acquisition

The study protocol for MEG, MRI, and DWI data was approved by the HUS ethical committee (**HUS/3043/2021**, 27.4.2022), written informed consent was obtained from each participant prior to the experiment, and all research was carried out according to the Declaration of Helsinki. The details of MEG and DWI data recording processing can be found in the Supplementary Materials.

### Model fitting

To fit the model simultaneously to both the synchronization and criticality metrics, we implemented a multi-objective gradient descent fitting algorithm. This approach allowed us to iteratively adjust model parameters to balance convergence to these two metrics. Since PLV exhibited a monotonically increasing behavior, we defined it as a sigmoid function (*S*(*P*)) where (*P*) were optimizable model parameters such as local or global coupling. The gradient was calculated based on the difference between the simulated and target FC matrices, denoted *loss*, while the sign of difference also showed the direction of the gradient:

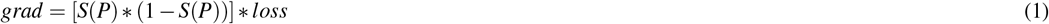

where *S* was a sigmoid function, *P* its parameter (e.g. global or local control parameter) and *loss* the element-wise difference between the target and simulated FC matrices.

The critical dynamics is characterized by the peak in the DFA curve as a function of the control parameter. Therefore, to approximate this relationship, we used the Lorentzian function (*g*(*P*)) due to its prominent peak and long tails. This allowed for the use of its derivative in gradient descent fitting, as shown in Equation 2:

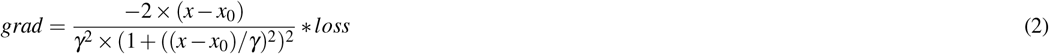

where the Lorentzian width parameter *γ* was equal to 0.5 for fitting neuroimaging data and 0.25 for model data and *x*_0_ corresponded to the average value of the parameters.

We used the multi-objective gradient descent fitting approach^73^ to integrate gradients from two objective functions. Since the control parameters influenced both criticality and phase synchrony, we computed the total gradient as a sum of DFA and PLV gradients. Because the DFA function is non-monotonic, we used the sign of the PLV gradient to determine the direction of the update (Figure 2C). Each gradient vector was normalized using its norm to ensure balanced contributions from both terms:

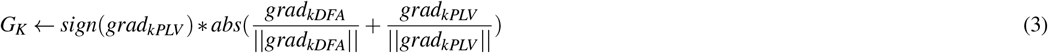

We applied L1 regularization with *λ* = 0.001 to avoid overfitting. In addition, we incorporated the Adam Optimizer for adaptive learning. The algorithm combining all of these steps can be found in the Supplementary Materials.

### Statistics

We used the Pearson correlation coefficient to evaluate the relationship between the symptom severity and both the observables and model parameters. Statistical significance was assessed using a two-sided t-test. We also performed linear regression to obtain the R-value, which quantified the association between the symptom severity and the average value of observable or parameter that showed a significant correlation. To estimate confidence intervals for the correlations between SDS symptom severity and the MEG observables or model parameters, we applied both jackknife resampling and bootstrap methods.

## Data and software availability

The MEG and DWI datasets generated and analyzed in the current study are not publicly available due to patient privacy limitations set forth in the Ethical Committee approval along with General Data Protection Regulation (GDPR) constraints. The model code are deposited in the publicly available repository (https://github.com/palvalab/hierarchical_kuramoto_zero) and the fitting code will be open-sourced following publication.

## Acknowledgments

The data preprocessing were performed using computer resources within the Aalto University School of Science “Science-IT” project. The simulations were performed on the LUMI supercomputer. This study was supported by the Academy of Finland (J.M.P, project number: 296304), by the Juselius Foundation (S.P and J.M.P, project number: 240156), and by Business Finland (J.M.P, project number: 1670/31/2024). Work on “PlaStim: Plasticity Stimulation in the Treatment of Anhedonia” was supported by Wellcome Leap as part of the Multi Channel Psych Program to S.P. and J.M.P.

