## Supplementary Material for "Mitigating activity mixing with personalized whole-brain modeling"

#### Supplementary Figures

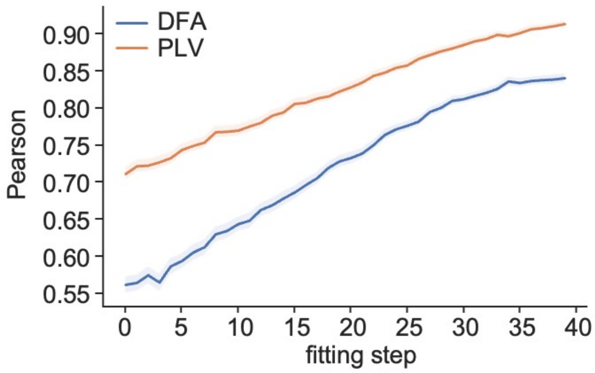


**Figure 1.** Increase in correlations between target and fitted observables over fitting steps.

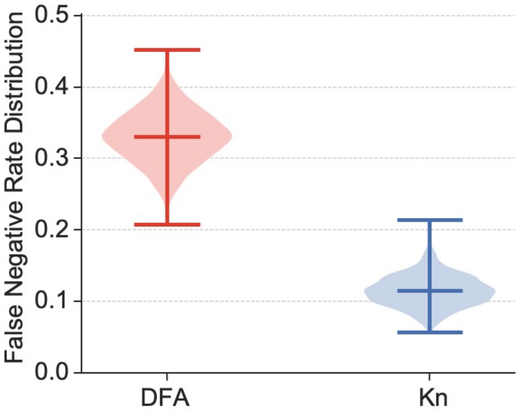


**Figure 2.** Bootstrapped False Negative Rate Distribution for the model DFA observables and fitted local coupling parameters.

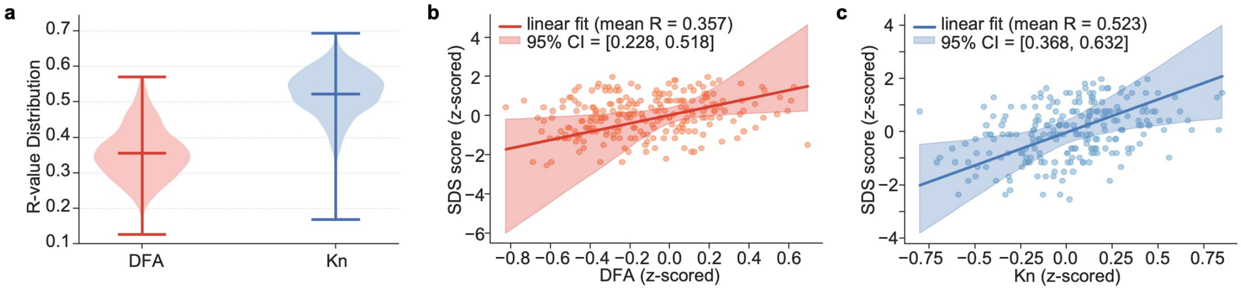


**Figure 3.** **a** Bootstrapped R-value for nodes and parcels significantly correlated with SDS symptoms for DFA observables and fitted local coupling parameters. **b** Scatter plot showing the relationship between z-scored SDS symptom values and z-scored DFA values from significantly correlated parcels, with linear regression. **c** Scatter plot of z-scored SDS symptom values versus z-scored values of significantly correlated nodes of local coupling parameters, with linear regression.

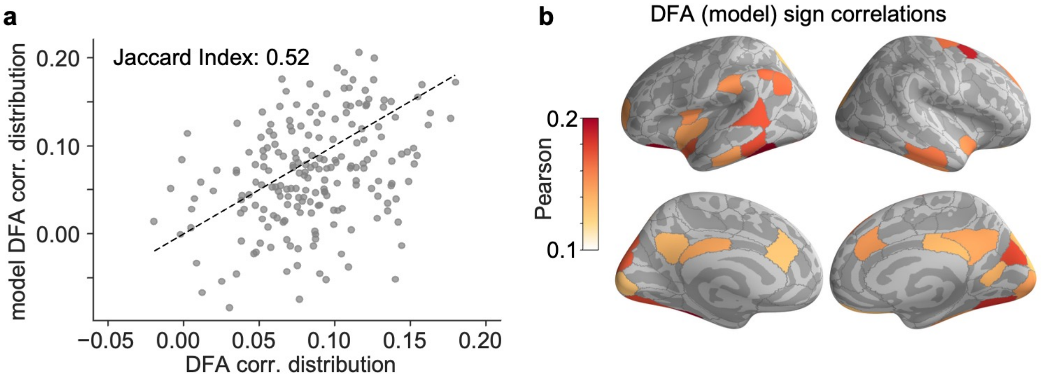


**Figure 4.** **a** Scatter plot of correlation values between SDS symptom values and patients' DFA against correlation values between SDS symptom values and DFA of the fitted model. **b** Anatomical distribution of significant Pearson coefficients between DFA of fitted model and SDS symptom across model nodes at 11Hz.

### MEG data preprocessing

Resting-state MEG data was recorded from all subjects over a 15-minute period using a 306-channel MEG system (Megin Oy, Helsinki) at two sites: the BioMag Laboratory at Helsinki University Hospital in Helsinki, Finland and the MEG Core at Aalto University in Espoo, Finland. Participants were instructed to keep their eyes open and focus on a cross at the center of a screen in front of them. Additionally, bipolar horizontal and vertical electrooculography (EOG) and bipolar electromyography (EMG) were recorded.

We used the MNE software (https://mne.tools/stable/index.html) for data preprocessing and source modeling. To suppress extracranial noise from MEG sensors, interpolate bad channels, and compensate head movements, we applied Temporal Signal Space Separation (tSSS) using the MaxFilter software (Elekta Neuromag Ltd., Finland). We performed Independent Component Analysis (ICA) to identify and remove artifact components linked to ocular activity using EOG signals, cardiac activity using the magnetometer signal as a reference, and muscle activity.

We conducted volumetric segmentation, surface reconstruction, cortical flattening, and parcellation of MRI data using FreeSurfer 7.3.2 (https://surfer.nmr.mgh.harvard.edu/). We constructed a surface-based source space with 5 mm spacing using a single-layer (inner skull) symmetric Boundary Element Method (BEM) for the forward operator. Noise covariance matrices were calculated from preprocessed data filtered between 151–249 Hz. To reduce spurious connections from source leakage, we estimated vertex fidelity and generated fidelity-weighted inverse operators. We aggregated inverse-transformed source time series into parcel time series optimizing source reconstruction accuracy[1,2]. We mapped the source time series to 200 parcels using the Schaefer atlas[3].

We obtained a narrowband representation of MEG signals using complex Morlet wavelets[4]. To suppress line noise at 50 Hz and its harmonics, we applied a notch filter with 1 Hz transition bandwidths.

### DWI data preprocessing

We preprocessed diffusion-weighted imaging (DWI) data, including denoising and motion correction, using the MRtrix3 3.0.1 Toolbox[5] and FMRIB Software Library (FSL) 6.0.5[6]. To minimize biases or artifacts from image acquisition or processing, we used the Advanced Normalization Tools (ANTs) 2.3.5[7]. For T1-weighted anatomical MRI preprocessing, head model creation, and cortical surface reconstruction, we used FreeSurfer 7.3.2 (https://surfer.nmr.mgh.harvard.edu/,[8]).

We applied Constrained Spherical Deconvolution (CSD)[9] to estimate fiber orientation distributions (FODs) within each voxel for the accurate decomposition of the diffusion signal into individual fiber orientations. To improve fiber tracking accuracy, we applied Anatomically Constrained Tractography (ACT,[10]), incorporating anatomical information from five-tissue-type (5TT) segmentation. The resulting FODs provided detailed maps of white matter tracts. Tractograms, generated from 20 million seed points, were refined using Spherical informed filtering of tractograms approach (SIFT2,[11]) to reduce reconstruction biases.

We coregistered T1-weighted anatomical MRI and DWI data using the SPM 12 package within MATLAB R2022a.

Structural connectomes (SCs) were constructed as symmetric edge-adjacency matrices, representing white matter connections between 200 parcels of the Schaefer atlas[3], aligned with Yeo 17 networks[12].

All preprocessing steps were performed on the Triton high-performance computing cluster running the CentOS Linux operating system at Aalto University.

### Fitting Algorithm

**Algorithm 1** Gradient Calculation

**Require:** $loss_{PLV}, loss_{DFA}, loss_{OCC}$
**Ensure:** $grad_K, grad_w$
**function** CALCULATEGRADIENTS($loss_{PLV}, loss_{DFA}, loss_{OCC}$)
  **if** regularization is True **then**
    Apply regularization to losses
    $loss_{DFA} \leftarrow loss_{DFA} + 1e^{-3} \times sign(loss_{DFA}) \times |K_n - 1|$
    $loss_{PLV} \leftarrow loss_{PLV} + 1e^{-3} \times sign(loss_{PLV}) \times |weights - 1|$
    $loss_{OCC} \leftarrow loss_{OCC} + 1e^{-3} \times sign(loss_{OCC}) \times |weights - 1|$
  **end if**
  $grad_{kPLV} \leftarrow sigmoid\_gradient(K_n) \times mean(loss_{PLV})$
  $grad_{wPLV} \leftarrow sigmoid\_gradient(weights) \times loss_{PLV}$
  $grad_{wOCC} \leftarrow lorentzian\_gradient(weights) \times loss_{OCC}$
  $grad_{kDFA} \leftarrow lorentzian\_gradient(K_n) \times loss_{DFA}$
  $grad_w \leftarrow grad_{wPLV}/||grad_{wPLV}|| + sign(grad_{wPLV}) \times |grad_{wOCC}|/||grad_{wOCC}||$
  $grad_K \leftarrow grad_{kPLV}/||grad_{kPLV}|| + sign(grad_{kPLV}) \times |grad_{kDFA}|/||grad_{kDFA}||$
  Apply masks and cutoff thresholds:
  $cutoff_{DFA} \leftarrow percentile(|loss_{DFA}|, 75)$
  $grad_K[|loss_{DFA}| < cutoff_{DFA}] \leftarrow 0$
  $grad_w \leftarrow grad_w \times |mask|$
  **return** $grad_K, grad_w$
**end function**
**function** SIGMOID_GRADIENT($x$)
  $f \leftarrow \frac{1}{1+e^{-x}}$
  **return** $f \times (1 - f)$
**end function**
**function** LORENTZIAN_GRADIENT($x, x_0, \gamma$)
  **return** $-2 \times (x - x_0)/(\gamma^2 \times (1 + ((x - x_0)/\gamma)^2)^2)$
**end function**
**Require:** $target_{PLV}, target_{DFA}, target_{OCC}$
**Ensure:** $weights, K_n$
**function** TUNEPARAMETERS($connectome, parameters, targets, mask, regularization, n\_iters, lr$)
  Initialize: $weights \leftarrow 1$, $K_n \leftarrow 1$
  **for** $i = 1$ to $n\_iters$ **do**
    Simulate model and compute losses:
    $loss_{PLV}, loss_{DFA}, loss_{OCC} \leftarrow ComputeLosses()$
    $grad_K, grad_w \leftarrow$ CALCULATEGRADIENTS($loss_{PLV}, loss_{DFA}, loss_{OCC}$)
    $update_K \leftarrow$ ADAM_OPTIMIZER($K_n, grad_K, lr$)
    $update_w \leftarrow$ ADAM_OPTIMIZER($weights, grad_w, lr$)
    $weights \leftarrow weights - update_w$
    $K_n \leftarrow K_n - update_K$
  **end for**
  **return** $weights, K_n$
**end function**
